# Cardiolipin synthesis in *Pseudomonas fluorescens* UM270 plays a relevant role in stimulating plant growth under salt stress

**DOI:** 10.1101/2022.10.19.512789

**Authors:** Daniel Rojas-Solis, Miguel A. Vences-Guzman, Christian Sohlenkamp, Gustavo Santoyo

## Abstract

Membrane cardiolipin (CL) phospholipids play a fundamental role in the adaptation of bacteria to various environmental conditions, including saline stress. Here, we constructed deletion mutants of two CL synthetase genes, *clsA* and *clsB*, in the rhizobacterium *Pseudomonas fluorescens* UM270, and evaluated their role in plant growth promotion under salt stress. *P. fluorescens* UM270 Δ*clsA* and Δ*clsB* mutants showed a significant reduction in CL synthesis compared to the UM270 wild-type strain (58% Δ*clsA* and 53% Δ*clsB*), and their growth rate was not affected, except when grown at 100 and 200 mM NaCl. Additionally, the root colonization capacity of both mutant strains was impaired compared with that of the wild type. Concomitant with the deletion of *clsA* and *clsB*, some physiological changes were observed in the UM270 Δ*clsA* and Δ*clsB* mutants, such as a reduction in indole acetic acid and biofilm production. By contrast, an increase in siderophore biosynthesis was observed. Further, inoculation of the UM270 wild-type strain in tomato plants (*Lycopersicon esculentum* Saladette) grown under salt stress conditions (100 and 200 mM NaCl) resulted in an increase in root and shoot length, chlorophyll content, and dry weight. On the contrary, when each of the mutants (Δ*clsA* and Δ*clsB*) were inoculated in tomato plants, a reduction in root length was observed when grown at 200 mM NaCl, but the shoot length, chlorophyll content, and total plant dry weight parameters were significantly reduced under normal or saline conditions (100 and 200 mM NaCl), compared to UM270 wild-type-inoculated plants. In conclusion, these results suggest that CL synthesis in *P. fluorescens* UM270 plays an important role in the promotion of tomato plant growth under normal conditions, but to a greater extent, under salt-stress conditions.

## 1. Introduction

Soil salinity is recognized as one of the main abiotic conditions that cause extensive losses to agricultural production worldwide (Flowers, 2004; Numan et al., 2018). Saline soil may contain salts, such as sulfate and chloride of calcium (Ca), magnesium (Mg), sodium (Na), and potassium (K) (Zaman et al., 2005), thus producing osmotic stress that affects water balance and ion homeostasis in plants. In addition, saline growing conditions alter the hormonal status of plants by disturbing transpiration, nutrient acquisition, and photosynthetic systems (Gupta and Huang, 2014; Ilangumaran and Smith, 2017). However, in many regions worldwide, saline soils represent the only arable areas; therefore, farmers require efficient strategies to cope with such stressful conditions to cultivate and generate good yields (Munns and Gilliham, 2015). One strategy is to use genetically modified plants (GMPs) or salt-resistant genotypes (SRGs) (Roy et al., 2014). However, not all regions in the world, such as in developing countries, have easy access to GMPs or SRGs. Therefore, a different way to help plants tolerate salt stress is to employ plant growth-promoting bacteria (PGPB) (Forni et al., 2017; Lugtenberg and Kamilova, 2009; Orozco-Mosqueda et al., 2022). PGPB can be easily isolated, characterized, and highly abundant in almost every soil or plant endosphere or associated rhizosphere (Fadiji et al., 2022a; Korenblum et al., 2020).

Species of the genus *Pseudomonas* are among the most studied PGPB because they have different mechanisms for promoting plant growth. For example, the production of phytohormones (e.g., indole-3-acetic acid, IAA), solubilization of nutrients in the rhizosphere, or the production of stimulating diffusible or volatile compounds may directly induce plant growth (Fadiji et al., 2022b). Other metabolites, such as hydrogen cyanide (HCN), phenazines, siderophores, lipopeptides, and 2,4-diacetylphloroglucinol (DAPG), as well as hydrolytic enzymes, such as proteases, cellulases, chitinases, and β-glucanases, have been shown to be effective in reducing or eliminating plant diseases caused by diverse microorganisms. This remarkable diversity of compounds and enzymes is considered an indirect way to promote plant growth, as well as the ability of *Pseudomonas* spp. to induce systemic resistance (ISR) in plants (Raaijmakers et al., 2009; Weller, 2007).

Other mechanisms, such as the production of biofilms and the ability to colonize environments such as the rhizosphere in PGPB, are also important to occupy spaces, thus restricting access to nutrients to potential pathogens, as well as to exercise direct mechanisms to promote plant growth in such microecosystems (Hernández-Salmerón et al., 2017; Rojas-Solis et al., 2016). However, under saline conditions in soils, bacteria must keep their membranes stable and, in turn, allow fluids and exchange of nutrients and metabolites, and ideally, keep PGP mechanisms active (Rojas-Solis et al., 2020; Soltani et al., 2005). Thus, bacteria use specific mechanisms to adapt to environmental conditions (including salt stress), such as adjusting their membrane phospholipid (PL) composition (Kondakova et al., 2015; Ramos et al., 1997)). *Pseudomonas* spp. possesses four major PL types: phosphatidylethanolamine (PE), phosphatidylglycerol (PG), phosphatidylcholine (PC), and cardiolipin (CL) (Geiger et al., 2013; Kondakova et al., 2015). CL is synthetized by cardiolipin synthase (*cls*), which catalyses the condensation of two PG molecules to yield CL and glycerol (Bernal et al., 2007; Von Wallbrunn et al., 2002). CL is a phospholipid that plays an important role in the adaptation of the cell membrane to saline stress (De Leo et al., 2009; Murínová and Dercová, 2014; Romantsov et al., 2007). For example, López et al., (2006) reported that a *Bacillus subtilis* strain deficient in CL production (*clsA* mutant) was impaired when grown in elevated NaCl concentrations. Another major membrane lipid, PE, has been shown to be important for the establishment of nitrogen-fixing root nodule symbiosis in *Sinorhizobium meliloti* (Vences-Guzmán et al., 2008). In another study, the bacterial strain L115 promoted the growth of peanuts (*Arachis hypogaea*) and tolerated high growth temperature and salinity by modifying the degree of fatty acid unsaturation and increasing phosphatidylcholine levels (Paulucci et al., 2015). To our knowledge, even though it is known that membrane lipid components are important for survival under osmotic stress, a specific role has not been associated with plant growth-promoting activities or coding genes for membrane components in any non-symbiotic bacteria. Therefore, here, we report the generation of two mutants in the synthesis of cardiolipin to determine the role of this important membrane phospholipid in plant growth-promoting activities in the rhizosphere strain *P. fluorescens* UM270 under normal and salt-stress conditions.

## 2. Materials and methods

### 2.1 Bacterial strains, media, and growth conditions

*P. fluorescens* UM270 strains were grown at 30 °C for 24 h in Luria Bertani (LB) medium, and routinely maintained at 4 °C. *Escherichia coli* strains were grown in LB medium at 37 °C. Antibiotics were added to the medium at the following concentrations when required (in micrograms per milliliter): 200 for neomycin and 100 for carbenicillin for *P. fluorescens*; 100 for carbenicillin, 50 for kanamycin, and 20 for tetracycline for *E. coli*.

### 2.2 Construction of the *clsA* and *clsB* deletion mutants in *P. fluorescens* UM270

The genome sequence of *P. fluorescens* UM270 was searched for the presence of genes encoding putative cardiolipin synthases (GenBank accession number: JXNZ00000000.1). Two sequences of *P. fluorescens* UM270, with accession numbers: KIQ59265.1 (*cls*A) and KIQ56391.1 (*clsB*), which encode the putative CL synthase genes *clsA* and *clsB*, were cloned, sequenced, and mutated. Oligonucleotide primers CLSA01 (5’-ACTGGAATTCCCTGCGCCGGGGTCAGGCAGACGCGAA-3’) and CLSA02 (5’-ACTGGGATCCCTGGGACGCGCGGGCGGTCAAAGCTTC-3’) were used for PCR amplification of the upstream region (∼1 kb) of the putative *clsA* gene from UM270, introducing EcoRI and BamHI sites into the PCR product. Similarly, primers CLSA03 (5’-ACTGGGATCCTAAGATCGTCTACGCCGCGTCCAGCCT-3’) and CLSA04 (5’-ACTGTCTAGAAACCTCGAGCAGTACTGCAAGACGCTG-3’) were used to amplify the downstream (∼1 kb) putative *clsA* gene from *P. fluorescens* by introducing BamHI and XbaI sites into the PCR product. After digestion with the respective enzymes, the PCR products were cloned as EcoRI/BamHI or BamHI/XbaI fragments into pUC19 to yield the plasmids (pUCD04) and (pUCD05), respectively. Then, the BamHI/XbaI fragment from (pUCD05) subcloned into (pUCD04) to yield (pUCD06). The plasmid (pUCD06) was digested with EcoRI and XbaI to subclone the regions usually flanking the *clsA* gene into the suicide vector pK18*mobsacB* (Schäfer et al., 1994) to yield pK18D01. After biparental mating using *E. coli* S17-1 as the mobilizing strain, pK18D01 was transferred into the UM270 wild-type strain. Transconjugants were selected on LB medium containing neomycin and nalidixic acid to select for single recombinants in the first step. The plasmid pK18*mobsacB* contains the *sacB* gene (Schäfer et al., 1994), which confers sucrose sensitivity in many bacteria. Growth of the single recombinants on 12.5 % (w/v) sucrose will select for double recombinants and the loss of the pK18*mobsacB* vector backbone from the bacterial genome. PCR analysis confirmed that the gene *clsA* was deleted. Similarly, the *clsB* deletion mutant was constructed. Oligonucleotide primers CLSB01 (5’-ACTGGAATTCCGTCGCGGCTGTCGTTCATCAGCAGCA-3’) and CLSB02 (5’-ACTGGGATCCGCGATTGTAGAAACGCAACTCCACCCC-3’) were used in a PCR to amplify approximately 1.0 kb of genomic DNA upstream of the putative *clsB* gene from UM270, introducing EcoR1 and BamHI sites into the PCR product. Similarly, primers CLSB03 (5’-ACTGGGATCCGGTGGACGACTGGGTCAGCATCGGCTC-3’) and CLSB04 (5’-ACTGTCTAGAGCAGTACTTCGATGACACCACCGGCCC-3’) were used to amplify about 1.0 kb of genomic DNA downstream of the putative *clsB* gene from *P. fluorescens* by introducing BamHI and XbaI sites into the PCR product. After digestion with the respective enzymes, the PCR products were cloned as EcoRI/BamHI or BamHI/XbaI fragments into pUC19 to yield the plasmids pUCD07 and pUCD08, respectively. The UM270 BamHI/HindIII fragment from pUCD08 was subcloned into pUCD07 to generate pUCD09. The plasmid pUCD09 was digested with EcoRI and XbaI to subclone the regions usually flanking *clsB* into the suicide vector pK18*mobsacB* (Schäfer et al., 1994) to yield pK18D02. Through biparental mating using *E. coli* S17-1 as a mobilizing strain, pK18D02 was introduced into the UM270 wild-type strain. Subsequent steps were performed as described above (similar to Δ*clsA*) to construct the *clsB* mutant. PCR and sequencing analyses confirmed that *clsA* and *clsB* were deleted from the UM270 genome, generating strains Δ*clsA* and Δ*clsB*. The bacterial strains and plasmids used in this study and their characteristics are listed in Table 1.

**Table 1.** Bacterial strains and plasmids use in this study.

| Strain or plasmid | Relevant characteristic (s) | Reference |
| --- | --- | --- |
| Strains | Wild type | Hernández-León et al., 2015 |
| *Pseudomonas fluorescens* UM270 |  |  |
| UM270 derivatives | UM270 carrying a deletion in *clsA* | This work |
| *ClsA*- |  |  |
| *ClsB*- | UM270 carrying a deletion in *clsB* | This work |
| *E. coli* | *recA1* _80 *lacZ*_M15; cloning strain |  |
| DH5α | | |
| S17-1 | *thi pro recA hsdR hsdM*_ RP4Tc::Mu Km::Tn7; Tpr Smr Spcr | Simon, R., et al 1983 |
| Plamids |  |  |
| pUC19 | Cloning vector, carbenicillin resistant | Yanisch-Perron, C., et at 1985 |
| pk18mobsacB | Suicide vector, kanamycin resistant | Schäfer, A., et al 1994 |
| pUCD04 | 1 kb fragment upstream of *clsA*, cloned as EcoR1/BamHI fragment in pUC19 | This work |
| pUCD06 | cloned as BamHI/Xba1 fragment in pUC19, 1 kb upstream and 1 kb downstream sequences | This work |
|  | flanking *clsA* cloned into pUC19 | This work |
| pK18D01 | Suicide vector for construction of mutant of *ClsA*- | This work |
| pUCD07 | 1 kb fragment upstream of *clsB*, cloned as EcoR1/BamHI fragment in pUC19 | This work |
| pUCD08 | 1 kb fragment downstream of *clsB*, cloned as BamHI/Xba1 fragment in pUC19 | This work |
| pUCD09 | 1 kb upstream and 1 kb downstream<br>sequences flanking *clsB* cloned into<br>pUC19 | This work |
| pK18D02 | Suicide vector for construction of<br>mutant of *ClsB*- | This work |

### 2.3 Determination of lipid composition of *P. fluorescens* UM270 and derivative mutants

The lipid composition of *P. fluorescens* UM270 wild-type and mutant strains was determined by labeling with [1-^14^C] acetate (Amersham Biosciences). Twenty-five milliliters of culture with 200 mM NaCl was adjusted to an optical density of 0.1. A 1 mL aliquot was transferred to a sterile tube, and 1 μCi of acetate was added. The cultures were incubated at 30 °C with shaking for 24 h. The cells of the larger cultures were harvested by centrifugation at 6000 rpm for 10 min at 4 °C. The pellets were washed with water and re-suspended in 100 μL water. Cells from the labelled cultures were centrifuged for 1 min at 14000 rpm, washed once, and resuspended in 100 μL water.

### 2.4 Quantitative analysis of lipid extracts

Lipids were extracted as previously reported, using a chloroform/methanol/water extraction (Sandoval-Calderón et al., 2015). Methanol: chloroform (375 µL, 2:1) was added to the suspended cells and the mixture was vortexed. Subsequently, 125 µL of water and 125 µL of chloroform were added to separate the lower and upper phases. The lower phase, which contained the lipids, was transferred to a new tube, washed once with water, dried under N_2_, and dissolved in a suitable volume of chloroform/methanol 1:1 (v/v). Aliquots of the lipid extracts were spotted onto HPTLC silica gel 60 plates (Merck, Pool, UK). Lipids were separated by two-dimensional TLC using chloroform/methanol/water (140:60:10, v/v/v) as the solvent for the first dimension and chloroform/methanol/glacial acetic acid (130:50:20, v/v/v) as the solvent for the second dimension. Unlabeled membrane lipids were visualized by iodine staining, and radioactive membrane lipids were visualized by exposure to autoradiography film (Kodak) or a Phosphor Imager screen (Amersham Biosciences). Individual lipids were quantified using the Image Quant software (Amersham Biosciences).

### 2.5 Survival experiments on salt stress

To evaluate the survival of UM270 and its mutants (Δ*clsA* and Δ*clsB*) under saline stress, strains were cultured in different concentrations of salt (0, 100, and 200 mM NaCl) in LB medium (supplemented with the respective NaCl concentrations). Growth was evaluated at an optical density (OD) of 590 nm (OD590) and determined starting off at an OD = 0.1; every 5 h of bacterial growth was monitored with a spectrophotometer.

### 2.6 Rhizosphere colonization experiments

To determine the rhizosphere or root colonization ability of the wild-type and mutant bacterial strains of *P. fluorescens* UM270, the method proposed by Scher et al. (1984) was employed, with some modifications (Rojas-Solis et al., 2016). Briefly, maize seeds were coated with each of the bacterial strains, wild type and mutants (Δ*clsA* andΔ*clsB*), and deposited in closed test tubes containing sterile raw soil and sand. Each seed was inoculated with approximately 10^4^ colony forming units (CFU) of each strain, and growth and colonization were further analyzed along the emerging radicle. The tubes were placed in a Percival growth chamber (Percival Scientific, Perry, IA, USA), with a photoperiod of 16 h light: 8 h dark at 22 °C for 30 days, and the CFU/g root of strains was measured.

### 2.7 Determination of the plant growth-promoting traits under salt stress

The IAA (indole-3-acetic acid) content was determined using the method described by Patten and Glick (2002), with some modifications. Briefly, 25 mL flasks were inoculated, supplemented with a graduated series of NaCl concentrations (0, 100, 200 mM NaCl) at 30 °C on a rotary shaker at 150 rpm. The cells were then collected by centrifugation at 10,000 *× g* for 15 min, and 2 mL of Salkowski reagent was added to the supernatant. The absorbance of the pink auxin complex was read at 540 nm using a UV– Vis spectrophotometer (JENWAY 7305). The calibration plot was constructed using dilutions of a standard total indole (Fluka, Switzerland) solution and uninoculated medium with the reagent as a control. The experiments were performed in triplicate.

Siderophore production was evaluated by growing wild-type and mutant strains on Chrome Azurol agar (CAS) medium supplemented (or not) with 0, 100, and 200 mM NaCl (Santoyo et al., 2019). All experiments were performed in triplicates.

The biofilm formation capacity of the UM270 wild-type and mutant derivative strains was analyzed following the protocol of Wei and Zhang (2006). Strains were grown in LB media, supplemented (or not) with salt (100 or 200 mM NaCl) to an O.D. of 1 and then diluted (1:1000) with fresh LB broth. A 0.5 mL diluted culture was transferred to an Eppendorf tube. Bacteria were incubated without agitation for 72 h at 30 °C, and the biofilm was quantified in this time. The biofilm was stained with 0.1% (w/v) crystal violet for 15 min at room temperature and then rinsed thoroughly with water to remove unattached cells and residual dye. Ethanol (95%) was used to solubilize the dye that had stained the biofilm cells. The absorbance of the solubilized dye (A_570_) was determined using a UV–Vis spectrophotometer (Jenway 7305). All experiments were independently performed in triplicate at least twice.

### 2.8 Evaluation of plant growth promotion by UM270, Δ*clsA*, and Δ*clsB* under normal (without salt) and salt stress conditions

Inoculating experiments of wild-type UM270, Δ*clsA*, and Δ*clsB* strains in tomato plants (*Lycopersicon esculentum* ‘Saladette’) under a greenhouse pot experiment were performed according to Rojas-Solís et al. (2018). Briefly, the experiments were carried out in pots (6 cm tall × 5 cm wide) with sterile peat moss (Sphaigne, Canada), with or without irrigation with salt solutions of 100 and 200 mM NaCl. First, tomato seeds were germinated *in vitro*, and after one week, seedlings of the same size were selected and transplanted into pots (one plant was left in each pot). Bacterial inoculants dissolved in sterile deionized water were applied every week after pot transplantation according to the experimental design, which also included treatments without bacterial inoculation. The concentration of the bacterial inoculants was adjusted such that their optical density at 600 nm was 1 (∼0.75–1×10^8^ UFC). Throughout the experiment, the plants were irrigated every third day with deionized water or saline solution, while the salt concentration was constantly controlled by measuring electrical conductivity (Field Scout. Mod. 2265FS). Each of the experimental treatments: control, control + NaCl, and control + each of the three strains (UM270 WT, Δ*clsA*, and Δ*clsB*) included 12 plants. The effect of each of the bacterial inoculants on root length, shoot length, total dry weight, and chlorophyll content was evaluated after five weeks of plant growth. Chlorophyll concentration was measured in three leaves of each plant, as previously reported (Orozco-Mosqueda et al., 2013).

## 3. Results

### 3.1 Effect of salinity on phospholipid metabolism in *P. fluorescens* UM270 WT and Δ*clsA* and Δ*clsB* mutants

The synthesis and percentage of diverse membrane phospholipids in the *P. fluorescens* UM270 strain under normal and NaCl stress growth conditions were evaluated. First, the wild-type UM270 strain significantly increased the production of CL at 200 mM NaCl, while a reduction in phosphatidylcholine (PC) was observed. Other phospholipids, such as phosphatidylethanolamine (PE) and phosphatidylglycerol (PG), remained unchanged when strain UM270 was grown in the presence of NaCl (Table 2). Interestingly, when Δ*clsA* and Δ*clsB* mutants were subjected to salt stress (200 mM NaCl), a significant reduction in CL synthesis was observed (58% Δ*clsA* and 53% Δ*clsB*), while an increase in the synthesis of PG and PC was noted (Table 4 and Figure 1). These results suggest that *P. fluorescens* UM270 increased CL synthesis under salt stress, and that *clsA* and *clsB* genes are important for the biosynthesis of such phospholipids (CL).

**Table 2.** Effect of salinity stress on the incorporation of [^14^C] acetate into phospholipids of *P. fluorescens* UM270.

| PL (%) | Growth conditions |  |  |
| --- | --- | --- | --- |
|  | 0mM NaCl | 100mM NaCl | 200mM NaCl |
| PE | 74.3 ± 5.7 | 69.8 ± 1.4 | 75.57 ± 2.8 |
| PG | 18.4 ± 4.0 | 22.8 ± 1.4 | 15.15 ± 0.8 |
| PC | 4.7 ± 0.7 | 4.4 ± 0.6 | 1.69 ± 0.7 * |
| CL | 2.5 ± 0.7 | 2.8 ± 0.6 | 7.59 ± 0.5 * |

PL: phospholipids, PE: phosphatidylethanolamine, PG: phosphatidylglycerol, PC: phosphatidylcholine and CL: cardiolipin. Values represent means ± SE of three independent experiments. \* indicate that the means differ significantly with respect to the control value (0 mM) according to Duncan's multiple range test ( $p < 0.05$ ).

**Fig. 1.**
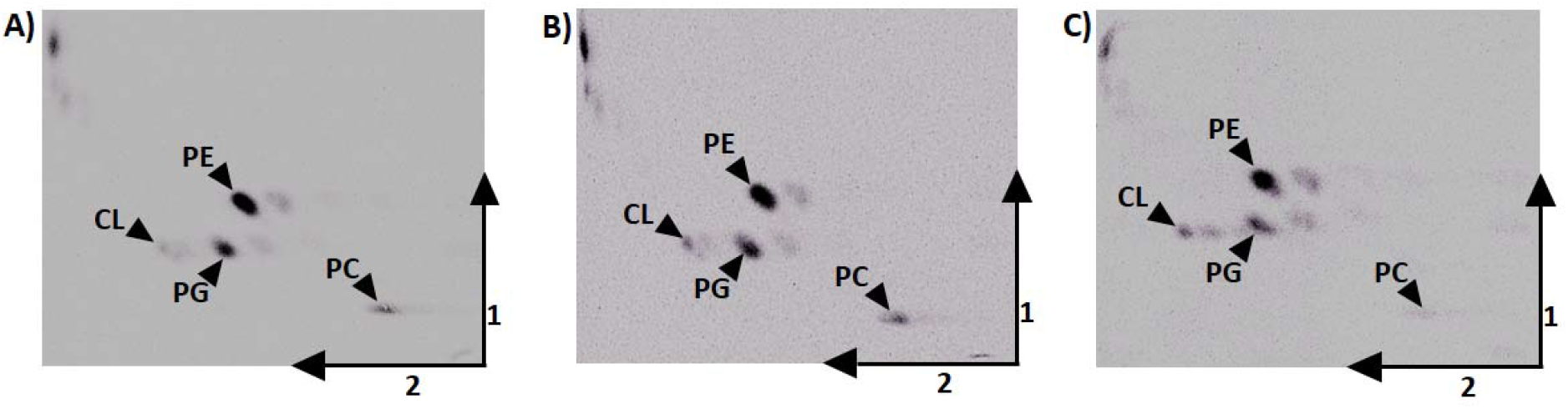
Membrane lipids of *P. fluorescens* UMS270 growth in LB medium supplemented (or not) with different salt concentrations: 0 mM (A), 100 mM (B), and 200 mM NaCl (C).

### 3.2 Growth of Δ*clsA* and Δ*clsB* mutants under normal and saline conditions

Given that CL phospholipids play an important role in cell membrane adaptation to different environmental stress conditions, we investigated the role of the *clsA* and *clsB* genes in salt stress growth (100 and 200 mM NaCl) in *P. fluorescens* UM270 and the Δ*clsA* and Δ*clsB* mutants (Figure 2). Deletion of the *clsA* or *clsB* genes did not affect growth in medium without salt stress; however, a similar and significant growth reduction was observed in the Δ*clsA* and Δ*clsB* mutants when grown at 100 and 200 mM NaCl, particularly during the exponential phase. This result suggests that *clsA* or *clsB*, which code for CL synthesis, are important for the growth of *P. fluorescens* UM270 under salt stress conditions.

**Fig. 2.**
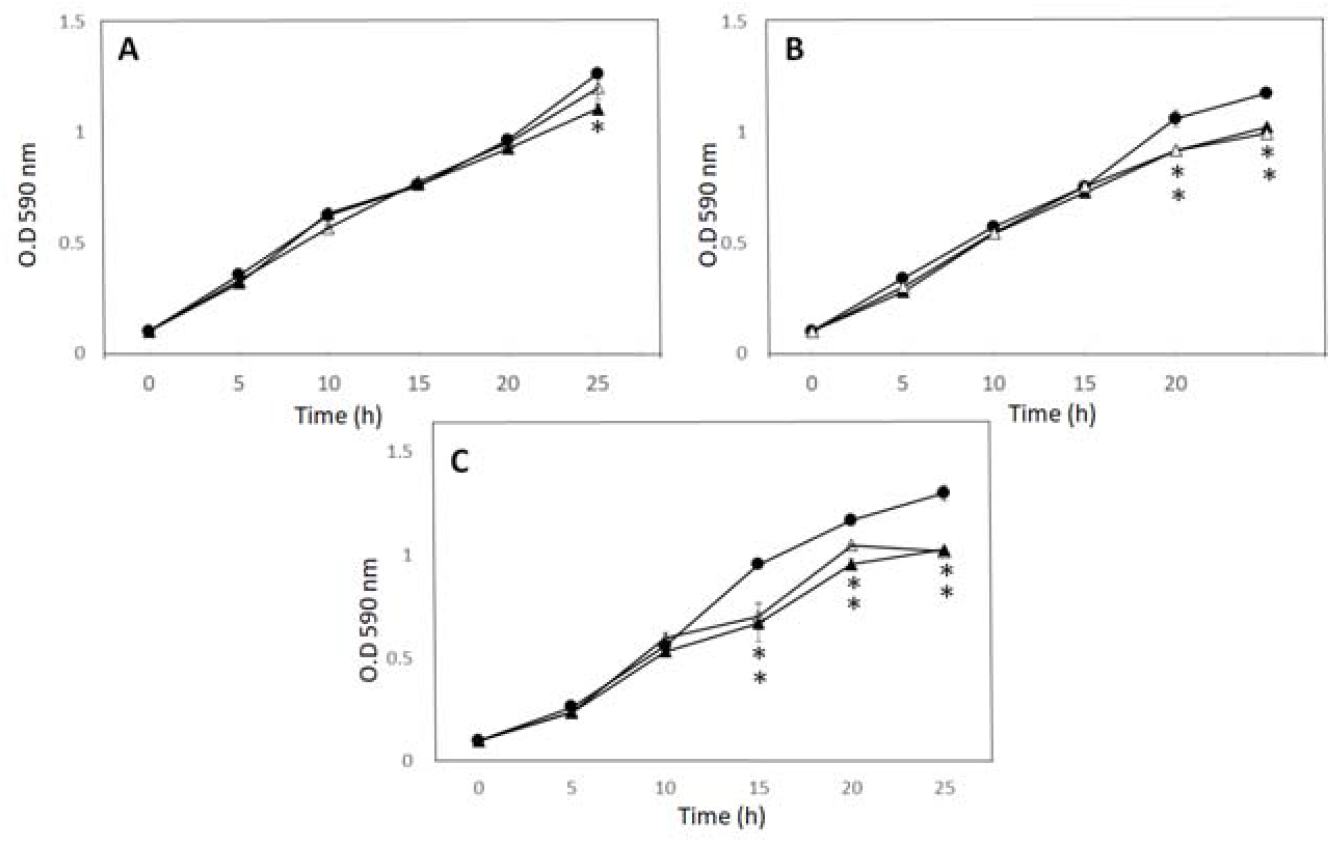
Growth of *P. fluorescens* mutants lacking *cls* is affected under saline conditions. *P. fluorescens* wild-type UM270 and mutants were grown in LB medium adjusted to 0 mM NaCl (A), 100 mM NaCl (B), or 200 mM NaCl (C). UM270 is represented by closed circles, Δ*clsA* by closed triangles, and Δ*clsB* by open triangles. Statistically significant growth inhibition was observed between treatment and control experiments marked by asterisks; Student’s *t*-test *P* < 0.05.

### 3.3 Rhizosphere colonization capacity of UM270 strains

The rhizosphere or root colonization capacity of strain UM270 has been previously evaluated (Rojas-Solis et al., 2016), showing similar results in these experiments. Table 3 shows the excellent colonization capacity of the wild-type strain to grow and colonize the roots of maize seedlings, as a significant increase in the number of CFU/g roots was recovered in the UM270 wild-type strain, but not in the Δ*clsA* and Δ*clsB* mutants.

**Table 3.**
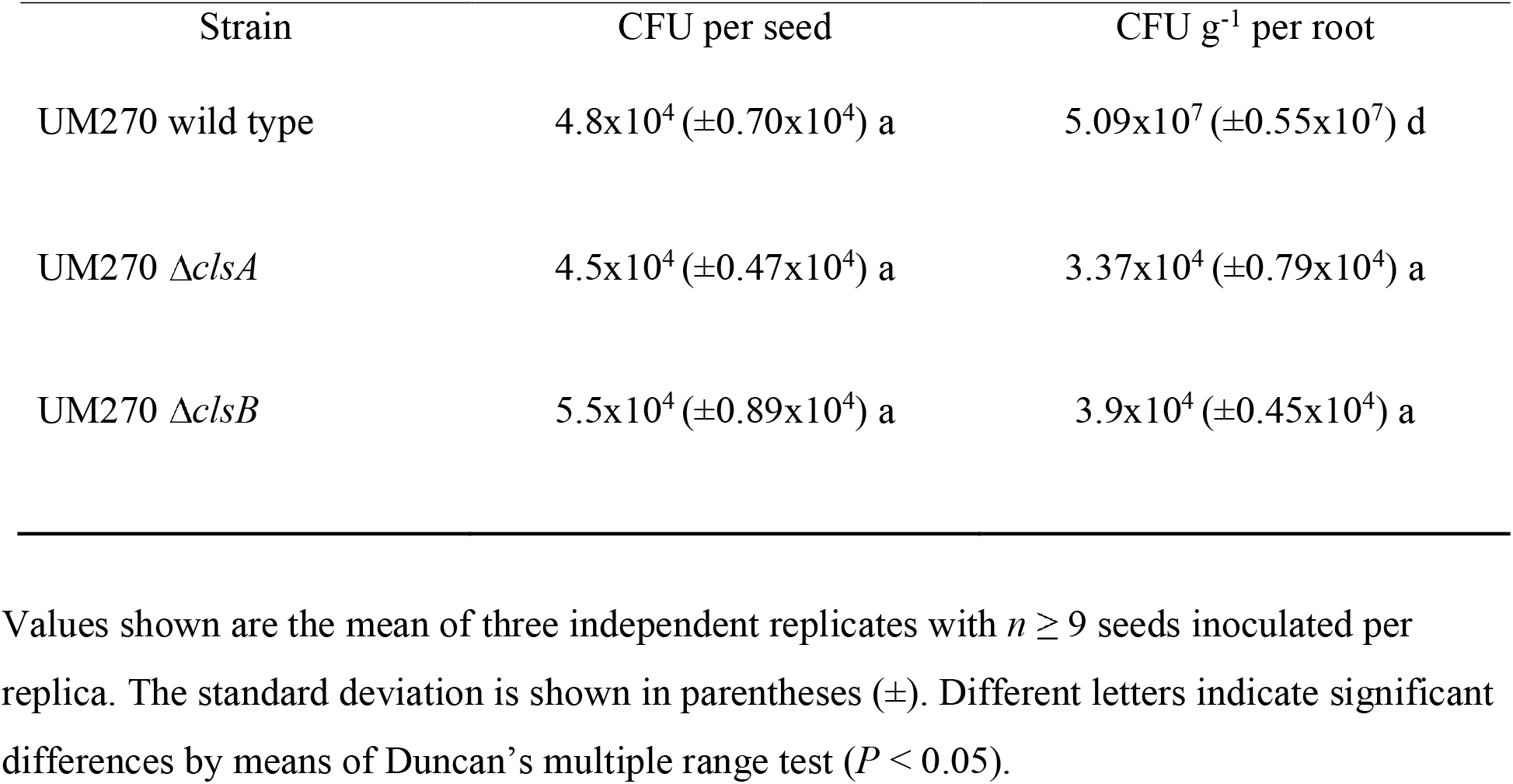
Rhizosphere colonization capacity of *Pseudomonas fluorescens* UM270 wild type and *cls* mutants.

| Strain | CFU per seed | CFU g<sup>-1</sup> per root |
| --- | --- | --- |
| UM270 wild type | 4.8x10<sup>4</sup> (±0.70x10<sup>4</sup>) a | 5.09x10<sup>7</sup> (±0.55x10<sup>7</sup>) d |
| UM270 Δ*clsA* | 4.5x10<sup>4</sup> (±0.47x10<sup>4</sup>) a | 3.37x10<sup>4</sup> (±0.79x10<sup>4</sup>) a |
| UM270 Δ*clsB* | 5.5x10<sup>4</sup> (±0.89x10<sup>4</sup>) a | 3.9x10<sup>4</sup> (±0.45x10<sup>4</sup>) a |

Values shown are the mean of three independent replicates with $n \geq 9$ seeds inoculated per replica. The standard deviation is shown in parentheses ( ± ). Different letters indicate significant differences by means of Duncan's multiple range test ( $P < 0.05$ ).

**Table 4.** Incorporation of (^14^C) acetate into phospholipids of *Pseudomonas fluorescens* UM270 wild-type and mutant strains under salt stress (200 mM NaCl).

| PL (%) | UM270 Wild-type | $\Delta clsA$ | $\Delta clsB$ |
| --- | --- | --- | --- |
| PE | 75.57 ± 2.8 | 74.9 ± 3.4 | 74.3 ± 3.2 |
| PG | 15.15 ± 0.8 | 19.5 ± 0.9 * | 19.2 ± 0.8 * |
| PC | 1.69 ± 0.7 * | 2.4 ± 0.4 | 3.0 ± 0.3 * |
| CL | 7.59 ± 0.5 | 3.2 ± 0.2 * | 3.5 ± 0.3 * |

PL: phospholipids, PE: phosphatidylethanolamine, PG: phosphatidylglycerol, PC: phosphatidylcholine and CL: cardiolipin. Values represent means ± SE of three independent experiments. \* indicate that the means differ significantly with respect to the control value (0 mM) according to Duncan's multiple range test ( $p < 0.05$ ).

### 3.4 Plant growth-promoting activities of *Pseudomonas fluorescens* UM270 and derivative Δ*clsA* and Δ*clsB* mutants

The deletion of the *clsA* or *clsB* gene in *P. fluorescens* UM270 resulted in important changes in the physiology of the mutant strains, particularly those related to some plant growth-promoting activities evaluated here. For example, both Δ*clsA* and Δ*clsB* mutants showed reduced indole acetic acid (IAA) production. This is, under control conditions (without salt), the UM270 wild-type strain produced 47.97 µg/mL of IAA, while the Δ*clsA* and Δ*clsB* mutants produced 20.45 µg/mL and 25.20 µg/mL of IAA, a 53.37 and 52.53% reduction, respectively. When IAA production in mutant strains was analyzed under salt stress, the negative effect was more pronounced (Table 5), reducing the IAA production to 2.78 µg/mL for Δ*clsA* and 2.63 µg/mL Δ*clsB* at 200 mM of salt.

**Table 5.** Summary of plant growth-promoting activities of *Pseudomonas fluorescens* UM270 and derivative Δ*clsA* and Δ*clsB* mutants in normal and saline conditions.

| Strain | NaCl (mM)<br>condition | IAA secreted<br>($\mu\text{g/ml}$) | Siderophore<br>production | Biofilm<br>Production |
| --- | --- | --- | --- | --- |
| UM270 | 0 | $47.97 \pm 3.6^a$ | $14.37 \pm 0.31^c$ | $0.038 \pm 0.0003^c$ |
| | 100 | $30.06 \pm 2.6^b$ | $14.62 \pm 0.42^c$ | $0.051 \pm 0.0003^{ab}$ |
| | 200 | $29.32 \pm 2.5^{bc}$ | $14.5 \pm 0.28^c$ | $0.053 \pm 0.004^b$ |
| $\Delta cIsA$ | 0 | $20.45 \pm 1.4^c$ | $18.87 \pm 0.68^b$ | $0.054 \pm 0.002^{ab}$ |
| | 100 | $8.07 \pm 1.9^d$ | $19.12 \pm 1.81^b$ | $0.064 \pm 0.01^b$ |
| | 200 | $2.78 \pm 1.4^d$ | $22.62 \pm 1.99^{ab}$ | $0.053 \pm 0.004^{ab}$ |
| $\Delta cIsB$ | 0 | $25.20 \pm 1.5^c$ | $21.75 \pm 1.12^{ab}$ | $0.060 \pm 0.003^{ab}$ |
| | 100 | $8.57 \pm 0.61^d$ | $19.75 \pm 1.73^{ab}$ | $0.044 \pm 0.002^{ab}$ |
| | 200 | $2.63 \pm 0.9^d$ | $23.37 \pm 1.28^a$ | $0.054 \pm 0.002^{ab}$ |

Siderophore production was evaluated as the halo zone diameter/colony diameter (mm). Biofilm production was measured at 570nm absorbance (D.O.). All data are mean values of three independent experiments, error bars indicate standard error. Different letters indicate that the means differ significantly according to Duncan's multiple range test ( $p < 0.05$ ). The statistical analysis was performed by PGP activity considering the three salinity conditions.

In contrast, siderophore synthesis was increased in both mutants (Table 5 and Supplementary Figure 1). Both Δ*clsA* and Δ*clsB* mutants presented a wider siderophore halo production either on normal CAS medium or CAS-Salt supplemented with 100 or 200 mM NaCl. Biofilm production was increased in the WT strain UM270 when subjected to salt stress conditions; however, for the Δ*clsA* and Δ*clsB* mutants, biofilm production under salt stress conditions did not present a significant increase compared to the control growth under normal conditions. It is worth mentioning that, even under normal conditions (no salt), both mutants produced more biofilm than the WT strain. This means that Δ*clsA* and Δ*clsB* mutants also exhibited increased biofilm production under both normal and salt stress conditions.

### 3.5 Tomato growth-promoting experiments

Greenhouse experiments were carried out to analyze the inoculation effect of the UM270 wild-type and mutants in CL on the growth of tomato plants under control and saline conditions (100 and 200 mM NaCl). The parameters evaluated in the tomato plants included root and shoot length, chlorophyll content, and total plant dry weight (biomass). First, inoculation with *P. fluorescens* UM270 WT increased the four parameters analyzed in tomato plants, namely, root and shoot length, chlorophyll content, and biomass content, either under normal or under stress caused by salt (100 or 200 mM NaCl). When the Δ*clsA* and Δ*clsB* mutants were inoculated in *Lycopersicon esculentum* grown in the greenhouse, a significant reduction in the root length was observed (only when growing in 200 mM NaCl), while the shoot length, chlorophyll content, and total plant dry weight were significantly reduced, either in normal or saline conditions (100 and 200 mM NaCl), compared to UM270 wild-type inoculated plants (Table 6). This result suggests that a mutation in either of the *cls* genes, which is important for the biosynthesis of membrane cardiolipin phospholipids, affects the natural plant growth-promoting traits of *P. fluorescens* UM270 (Figure 3).

**Table 6.**
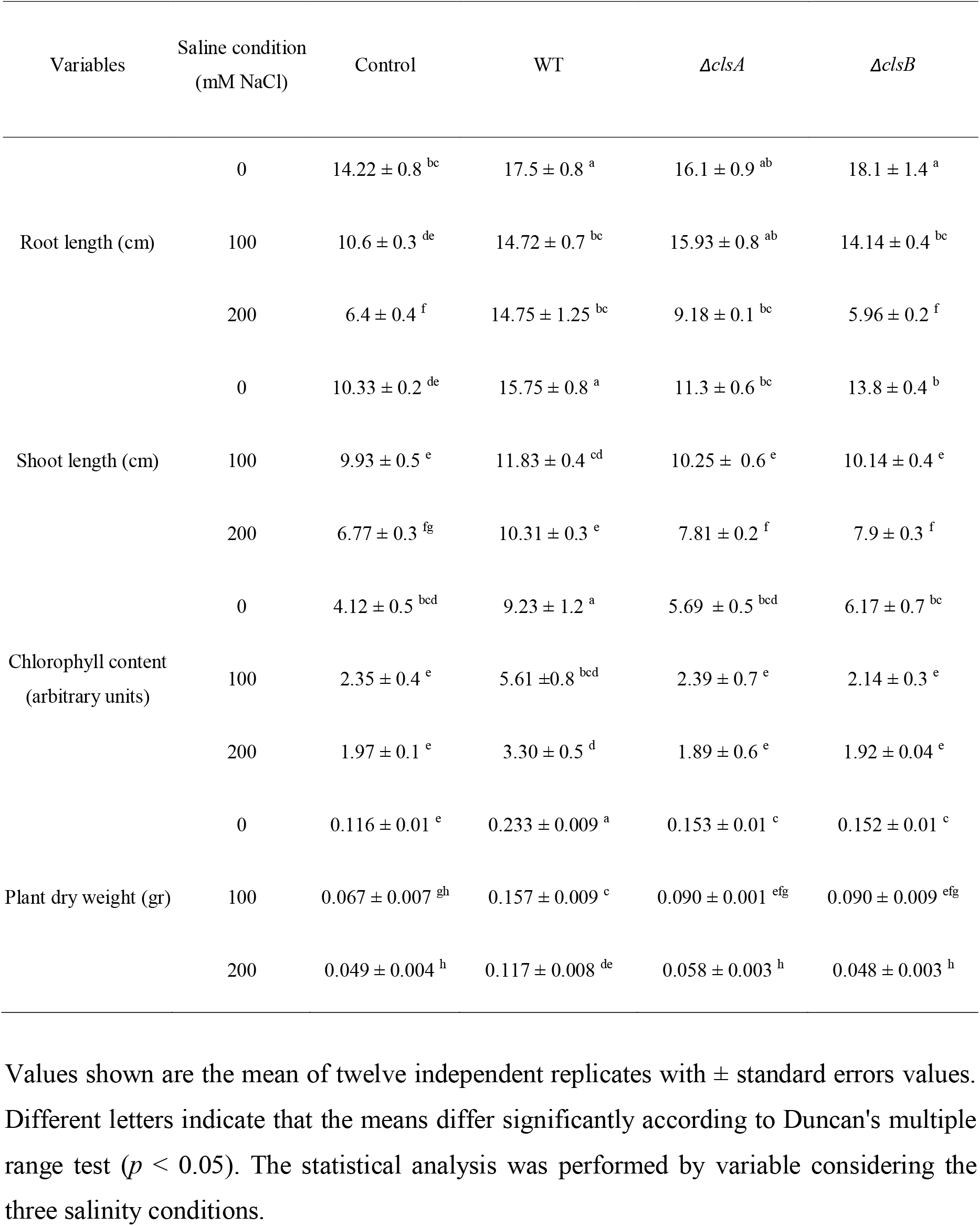
Tomato plants (*Lycopersicum esculentum* cv Saladette) growth promotion by inoculation of *Pseudomonas fluorescens* UM270 and mutant strains in normal conditions and saline stress.

| Variables | Saline condition<br>(mM NaCl) | Control | WT | $\Delta cIsA$ | $\Delta cIsB$ |
| --- | --- | --- | --- | --- | --- |
| Root length (cm) | 0 | 14.22 ± 0.8<sup>bc</sup> | 17.5 ± 0.8<sup>a</sup> | 16.1 ± 0.9<sup>ab</sup> | 18.1 ± 1.4<sup>a</sup> |
| | 100 | 10.6 ± 0.3<sup>de</sup> | 14.72 ± 0.7<sup>bc</sup> | 15.93 ± 0.8<sup>ab</sup> | 14.14 ± 0.4<sup>bc</sup> |
| | 200 | 6.4 ± 0.4<sup>f</sup> | 14.75 ± 1.25<sup>bc</sup> | 9.18 ± 0.1<sup>bc</sup> | 5.96 ± 0.2<sup>f</sup> |
| Shoot length (cm) | 0 | 10.33 ± 0.2<sup>de</sup> | 15.75 ± 0.8<sup>a</sup> | 11.3 ± 0.6<sup>bc</sup> | 13.8 ± 0.4<sup>b</sup> |
| | 100 | 9.93 ± 0.5<sup>e</sup> | 11.83 ± 0.4<sup>cd</sup> | 10.25 ± 0.6<sup>e</sup> | 10.14 ± 0.4<sup>e</sup> |
| | 200 | 6.77 ± 0.3<sup>fg</sup> | 10.31 ± 0.3<sup>e</sup> | 7.81 ± 0.2<sup>f</sup> | 7.9 ± 0.3<sup>f</sup> |
| Chlorophyll content<br>(arbitrary units) | 0 | 4.12 ± 0.5<sup>bcd</sup> | 9.23 ± 1.2<sup>a</sup> | 5.69 ± 0.5<sup>bcd</sup> | 6.17 ± 0.7<sup>bc</sup> |
| | 100 | 2.35 ± 0.4<sup>e</sup> | 5.61 ± 0.8<sup>bcd</sup> | 2.39 ± 0.7<sup>e</sup> | 2.14 ± 0.3<sup>e</sup> |
| | 200 | 1.97 ± 0.1<sup>e</sup> | 3.30 ± 0.5<sup>d</sup> | 1.89 ± 0.6<sup>e</sup> | 1.92 ± 0.04<sup>e</sup> |
| Plant dry weight (gr) | 0 | 0.116 ± 0.01<sup>e</sup> | 0.233 ± 0.009<sup>a</sup> | 0.153 ± 0.01<sup>c</sup> | 0.152 ± 0.01<sup>c</sup> |
| | 100 | 0.067 ± 0.007<sup>gh</sup> | 0.157 ± 0.009<sup>c</sup> | 0.090 ± 0.001<sup>efg</sup> | 0.090 ± 0.009<sup>efg</sup> |
| | 200 | 0.049 ± 0.004<sup>h</sup> | 0.117 ± 0.008<sup>de</sup> | 0.058 ± 0.003<sup>h</sup> | 0.048 ± 0.003<sup>h</sup> |

Values shown are the mean of twelve independent replicates with ± standard errors values. Different letters indicate that the means differ significantly according to Duncan's multiple range test ( $p < 0.05$ ). The statistical analysis was performed by variable considering the three salinity conditions.

**Fig. 3.**
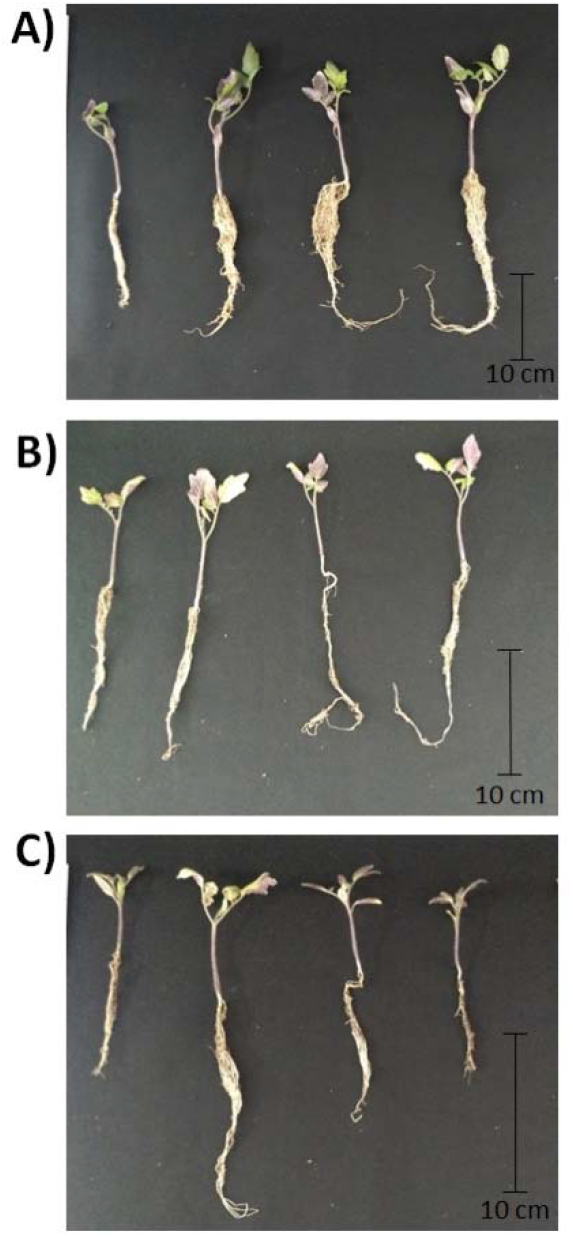
Composite picture of tomato (*L. esculentum* cv Saladette) plants inoculated with the indicated bacterial strains. Plants irrigated with water (A), plants irrigated with 100 mM NaCl (B), and plants irrigated with 200 mM NaCl (B). From left to right, plants were un-inoculated or inoculated with *P. fluorescens* wild-type UM270, Δ*clsA* mutant, and Δ*clsB* mutant. See the Section “Materials and Methods” for details of the experiment.

## 4. Discussion

The phospholipid cardiolipin (CL) plays an essential role in the adaptation of many bacterial species to environmental stresses including salinity (López et al., 2006; López et al., 2016; Romantsov et al., 2007). In the present study, we analyzed the role of reduced CL levels in response to saline stress in the plant growth-promoting bacterium *P. fluorescens* UM270 by deleting *clsA* and *clsB*. Each of the single mutations affected the production of the CL content (58% for Δ*clsA* and 53% for Δ*clsB*), indicating that both genes are involved in the CL synthesis in strain UM270. It is important to mention that both *clsA* and *clsB* are not part of an operon in the UM270 genome. A similar genetic array is found in other sister pseudomonads, such as *P. fluorescens* F113 (Miguel et al., 2012) and *P. putida*, which contain *clsA* and *clsB* as single genes expressed under the control of sigma factor (Bernal et al., 2007; Von Wallbrunn et al., 2002). When analyzing cell growth under saline conditions, the mutant strains showed only a slightly reduced delay in growth during the exponential phase. These results contrast with what has been reported, in which it was mentioned that CL deficiency causes delayed growth that occurs until the stationary phase in several bacterial species, such as *Agrobacterium tumefaciens* (Cronan, 2003). However, *P. fluorescens* operates two distinct cardiolipin synthases, similar to those in *A. tumefaciens* (*cls1* and *cls2* genes) (Czolkoss et al., 2016). López et al. (2006) reported that a *Bacillus subtilis* strain deficient in CL production was unable to grow at high salt concentrations. This result indicates that the presence of CL in the membranes is important for salinity adaptation; however, in this study, it was observed that a reduction in CL in the *P. fluorescens* UM270 membranes, although important for adaptation to salinity, is not essential for growth under these conditions. Several studies have proposed that the structure of CL, which consists of a quadruple-chained anionic amphiphile phospholipid composed of two 1,2-diacyl phosphatidate moieties esterified to the 1-and 3-hydroxyl groups of a single glycerol molecule, whose polar headgroup contains two phosphodiester moieties, confers a larger headgroup volume compared with other phospholipids, allowing the increase in CL to improve the structural integrity of the cell membrane under stressful conditions (Hoch, 1998; Lewis and McElhaney, 2009; Murínová and Dercová, 2014).

The ability of wild-type strain UM270 and cardiolipin mutants to maintain plant growth-promoting mechanisms under normal and salt stress conditions, including the production of indole-3-acetic acid (IAA), siderophore excretion, and biofilm production, was also evaluated. Such mechanisms are important for beneficial bacteria to promote plant growth (Hernández-León et al., 2015; Santoyo, 2021). The wild-type UM270 strain showed a significant reduction in IAA production, while siderophores were maintained and biofilm production was increased, as expected. Interestingly, both *cls* deletion mutants showed a reduction in IAA production compared to the mutant cells without salt stress. However, IAA production was reduced under salt stress in both the *cls* mutants. This result is contrary to what Mohamed and Gomaa (2012) reported that a *P. fluorescens* strain maintained its capacity to produce IAA under salt stress, which promoted the growth of radish plants (*Raphanus sativus*).

In contrast, siderophore and biofilm production were higher in both UM270 *cls* mutants when grown under saline conditions. For unknown reasons, the mutants suffered a physiological change, and siderophore production was also increased under normal conditions (see supplementary Figure 1). Unexpected physiological changes have also been observed in a 1-aminocyclopropane-1-carboxylate (ACC) deaminase deletion mutant of the endophytic plant growth-promoting bacterium *Burkholderia phytofirmans* PsJN, including an increase in indole acetic acid synthesis and a decrease in siderophore production (Sun et al., 2009). The authors suggested that a mutation in the nonessential gene (*acdS*) caused a change in the expression of stationary-phase sigma factor, but the reasons are still unknown. It would be important to go deeper into the analysis of the same factor, but also to carry out a transcriptional analysis that sheds light on other collateral changes that are created by mutating certain non-essential genes, such as *acdS* in *B. phytofirmans* PsJN or *clsA* and *clsB* in *P. fluorescens* UM270.

The production of biofilms in rhizobacteria is essential for the adaptation and colonization of this microenvironment. Rhizobacteria secrete a layer of extracellular polymeric substances that encapsulate cells and protect them from diverse environmental stresses (Lin et al., 2015). Here, biofilm formation did not present significant changes among the three conditions analyzed, and surprisingly, biofilm formation increased in the CL-reduced mutants in comparison with the wild-type strain in conditions without salt. These results differ from those reported by Lin et al. (2015), who reported that CL-deficient mutants of *Rhodobacter sphaeroides* generated ellipsoid-shaped cells that directly reduced biofilm formation.

Glycophyte plants, such as tomato plants (*Lycopersicon esculentum*), are forced to induce tolerance to salinity at salt concentrations as low as 10 mM NaCl (the salt tolerance grade also depends on the cultivar) to avoid adverse effects on growth and productivity (Bui and Henderson, 2003). Here, the plant growth stimulating capacity under normal and saline stress in tomato (*Lycopersicon esculentum* ‘Saladette’) plants exerted by *P. fluorescens* UM270 wild-type and Δ*clsA* and Δ*clsB* mutants was evaluated. As expected, saline conditions reduced the growth, chlorophyll content, and dry weight of plants, but the inoculation of the UM270 wild-type counteracted such stressful conditions. Plants inoculated with CL-reduced mutants did not show a recovery in the parameters evaluated, mainly in terms of chlorophyll content and biomass.

Under saline conditions, plants face osmotic and ionic stress and are forced to change their physiology to survive (Forni et al., 2017). Osmotic stress causes dehydration, whereas ionic stress leads to an excessive influx of Na ions, which can inhibit various basic physiological processes, including photosynthesis (Horie et al., 2012). Subsequently, the stress hormone ethylene increases and triggers other plant processes such as leaf yellowing, senescence, and premature death (Orozco-Mosqueda et al., 2020). To counteract these physiological changes, plants can interact with plant growth promoting bacteria, such as *P. fluorescens* UM270. Strain UM270 contains multiple plant growth-promoting mechanisms, such as ACC deaminase production, that reduce ethylene levels in stressed plants (Glick, 2014; Hernández-León et al., 2015). Additionally, UM270 produces siderophores, IAA, and biofilm, among other mechanisms that might act under such stressful conditions to maintain the growth promotion of tomato plants. However, the colonization capacity of *clsA* and *clsB* mutants is impaired, and therefore, UM270 may not have a beneficial effect on the plant because the number of colony-forming units necessary to exert such a beneficial effect on the plant is not reached in the rhizosphere of tomato plants (Santoyo et al., 2021). Few studies have evaluated changes in membrane phospholipids in bacteria and how these modifications alter the plant-bacteria interactions. For example, Vences-Guzmán et al. (2008) reported that a phosphatidylethanolamine (PE)-deficient *Sinorhizobium meliloti* strain is affected by its ability to nodulate alfalfa plants. The few nodules formed by the PE mutant strain were unable to fix nitrogen, while the leaves of the plants were yellowish, indicating nitrogen starvation symptoms. The authors concluded that PE and changes in lipid composition were important for the establishment of nitrogen-fixing root nodule symbiosis. Here, we evaluated the mutation in a different phospholipid, such as CL, since this phospholipid proportion was increased in the wild-type UM270 strain under salt stress conditions (mainly at 200 mM NaCl). It is important to evaluate its participation during interactions with plants under salt stress by mutating the genes responsible for CL biosynthesis. In conclusion, the CL membrane phospholipid, produced by the bacterium *P. fluorescens* UM270, might be important for root colonization and survival in the rhizosphere under conditions of saline stress to be able to exert its growth-promoting activities during interaction with tomato plants.

## Acknowledgement

This study was funded by Consejo Nacional de Ciencia y Tecnología, México (Grant number: A1-S15956) and CIC-UMSNH (2019–2020). DR-S thanks the PhD scholarship from Consejo Nacional de Ciencia y Tecnología, México. We thank Carlos Alberto Urtis-Flores for excellent technical assistance.

## Figure Legends

**Suppl. Fig. 1.**
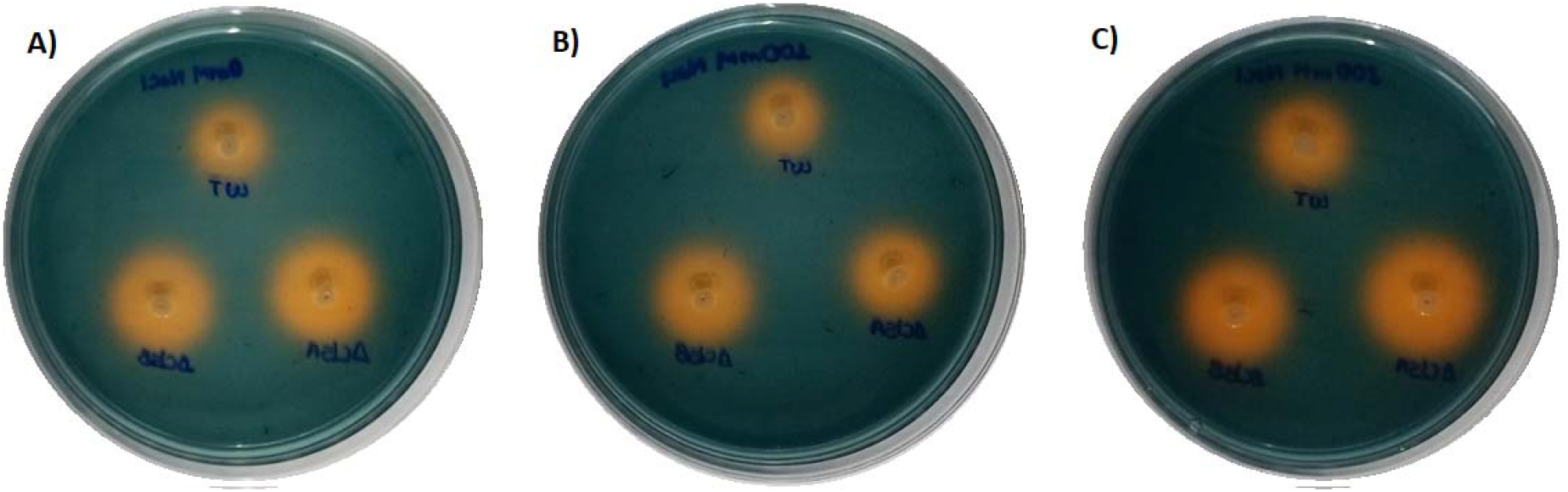
Effect of NaCl on siderophore production in *P. fluorescens* UM270 and *cls* mutants grown on CAS agar. In addition, WT *P. fluorescens* UM270, and below, from left to right, the Δ*clsB* and Δ*clsA* mutants.

